# New insights into 4,000 years of resource economy across Greenland using ancient DNA

**DOI:** 10.1101/2022.02.23.480846

**Authors:** Frederik V. Seersholm, Hans Harmsen, Anne Birgitte Gotfredsen, Christian Koch Madsen, Jens Fog Jensen, Jørgen Hollesen, Morten Meldgaard, Michael Bunce, Anders J. Hansen

## Abstract

The success and failure of past cultures across the Arctic was tightly coupled to the ability of past people to exploit the full range of resources available to them, and to adapt to fluctuations in resource availability. There is substantial evidence for the hunting of birds, caribou and a wide range of marine mammals in pre-historic Greenland from bone remains preserved in ancient middens. However, the extent to which these communities relied on marine resources such as fish and large cetaceans is understudied because of the taphonomic processes and bias that affect how these taxa present themselves in the archaeological record. To address this, we analyse DNA from bulk bone samples from 12 archaeological sites across Greenland dating from Paleo-Inuit to Neo-Inuit periods. Using a combination of metabarcoding and shotgun metagenomics we identify an assemblage of 43 species consisting of birds, fish, and both marine and terrestrial mammals. We find genetic evidence of five different whale species, of which the bowhead whale (*Balaena mysticetus*) was the most commonly detected. Furthermore, we detect nine fish species, of which four have not previously been identified in any of the studied sites. Lastly, we identify a novel haplotype in caribou (*Rangifer tarandus*) at the 3,000-year-old site Itinnera, suggesting the presence of a distinct lineage of (now extinct) dwarfed caribou that colonised Greenland after the last ice age 9,000 years ago. Collectively, these findings provide a rare insight into whaling and fishing practices in Greenland and demonstrate that prehistoric Greenlandic communities had the social and technological capacity to target the largest whales available in the waters around them.

## Introduction

The extreme cold temperatures north of the Arctic Circle have fostered an environment with a barren terrestrial ecosystem of low floral and faunal diversity. Despite these conditions, people have colonized the eastern Arctic in at least three distinct main migration waves^1^. The first migration into Greenland was formed by the Saqqaq people who settled along the coasts ~4,500 years ago^2,3^ but disappeared again some 1,700 years later, replaced by the Greenlandic Dorset culture (c. 800 BC – 1 AD)^4^ and, after a hiatus, the Late Dorset (.c 800-1300 AD). At approximately 985 AD, the descendants of the Vikings, the Norse, arrived in South Greenland^5^. They established two settlements, the Eastern and Western Settlement, and survived for approximately 500 years until the middle of the 15^th^ century, after which they disappeared as well^6^. The last group of people to settle in Greenland was the Thule culture who by current estimates arrive sometime in the 13^th^ century AD^7^. With roots in eastern Siberia, the Thule people were already adapted to a life in the Arctic, and they are the only people that have persisted in Greenland until the present day^8^. However, the Thule culture came under increasing European influence from the late 16th century; first by whalers and then by Danish-Norwegian colonial rule beginning in AD 1721. These increasing interactions had profound impacts on the Inuit, especially in regard to settlement and subsistence patterns (e.g. access to new technologies such as rifles, nets, iron hooks and harpoon heads).^9,10^

Compared to what is known about the earlier Paleo-Inuit cultures in the Arctic, the Thule culture stands out because of their sophisticated technology, such as the dogsled^11^, the kayak and the large skin boat^3^ or *umiaq*, that allowed them to travel long distances and hunt larger marine mammals. Accordingly, the Thule people were able to exploit the full range of subsistence animals available in Greenland, from small birds and fish to large whales. Although less archaeological material exists from the Paleo-Inuit cultures, the apparent absence of an *umiaq* sized vessel and the near absence of large harpoon heads suggests that the Paleo-Inuit may have had a more limited range of subsistence animals. For example, walrus and large whales are rarely found in Paleo-Inuit middens, while they are relatively abundant in archaeological sites from the Thule culture^12^. This apparent difference in subsistence strategies could help explain the disappearance of the Paleo-Inuit and the success of the Thule culture. Likewise, the changing climate during the Little Ice Age (1400-1900 AD) has been put forward as one of several factors contributing to the demise of the Norse colonization in Greenland around AD 1450^13^.

It is generally accepted that the survival of past cultures in the Arctic was heavily dependent on their ability to adapt to changing environmental conditions with e.g. new subsistence strategies when required^14^. Testing that hypothesis, however, has proven difficult for some groups of species as the quantifiable data on past subsistence practices relies heavily on preserved faunal remains found in archaeological deposits^2^, which are subject to various taphonomic processes^15^. The importance of whaling, for example, is notoriously difficult to pinpoint based on bone fragments excavated from middens as the meat and blubber from a whale is often exploited without bringing bones back to the settlement^16,17^. In most midden layers, the majority of large cetacean remains consist of artefacts or worked fragments of whalebone and (where preservation conditions are permitting) baleen. The fragmentary nature of large whale remains in middens further hampers species identification. Furthermore, while fish must be assumed to have contributed to the diet in Greenland, it is difficult to quantify as fish bones are generally small, cryptic, fragile and easily degrade under certain conditions^18^.

To illuminate aspects of the past Greenlandic resource exploitation that might be missed by traditional zoo-archaeological methods, we analysed faunal remains from across Greenland using a genetics approach. We applied bulk bone metabarcoding (BBM)^19^ on 2,500 small unidentifiable (usually fragmented) fossil bones, excavated from 12 distinct archaeological sites, representing the Paleo-Inuit (Saqqaq; n=4), Norse (n=2) and Neo-Inuit (Thule) culture (n=6).

## Results & Discussion

We collected 25 bulk bone samples of 2×50 bone fragments (2,500 bone fragments in total) from 12 archaeological sites across Greenland (Figure 1; Supplementary Material, Table 1 and Table 2). We analysed the samples using four metabarcoding assays targeting the *12S* or *16S* genes of mammals, vertebrates, birds and fish, respectively (Supplementary Material, Table 3). In total, we sequenced 22,907,439 reads, representing 298 amplicon sequence variants (ASVs; Supplementary Material, Table 4, Table 5, and Table 6). Of these, 238 ASVs (72%) could be confidently assigned to a taxon, which after filtering (Methods) represent 42 species (20 mammals, 13 birds and 9 fish; Supplementary Material, Table 7 and Table 8). Of the twelve archaeological sites analysed, eleven yielded endogenous DNA. Accordingly, all down-stream analyses were performed on these 11 sites.

**Figure 1.**
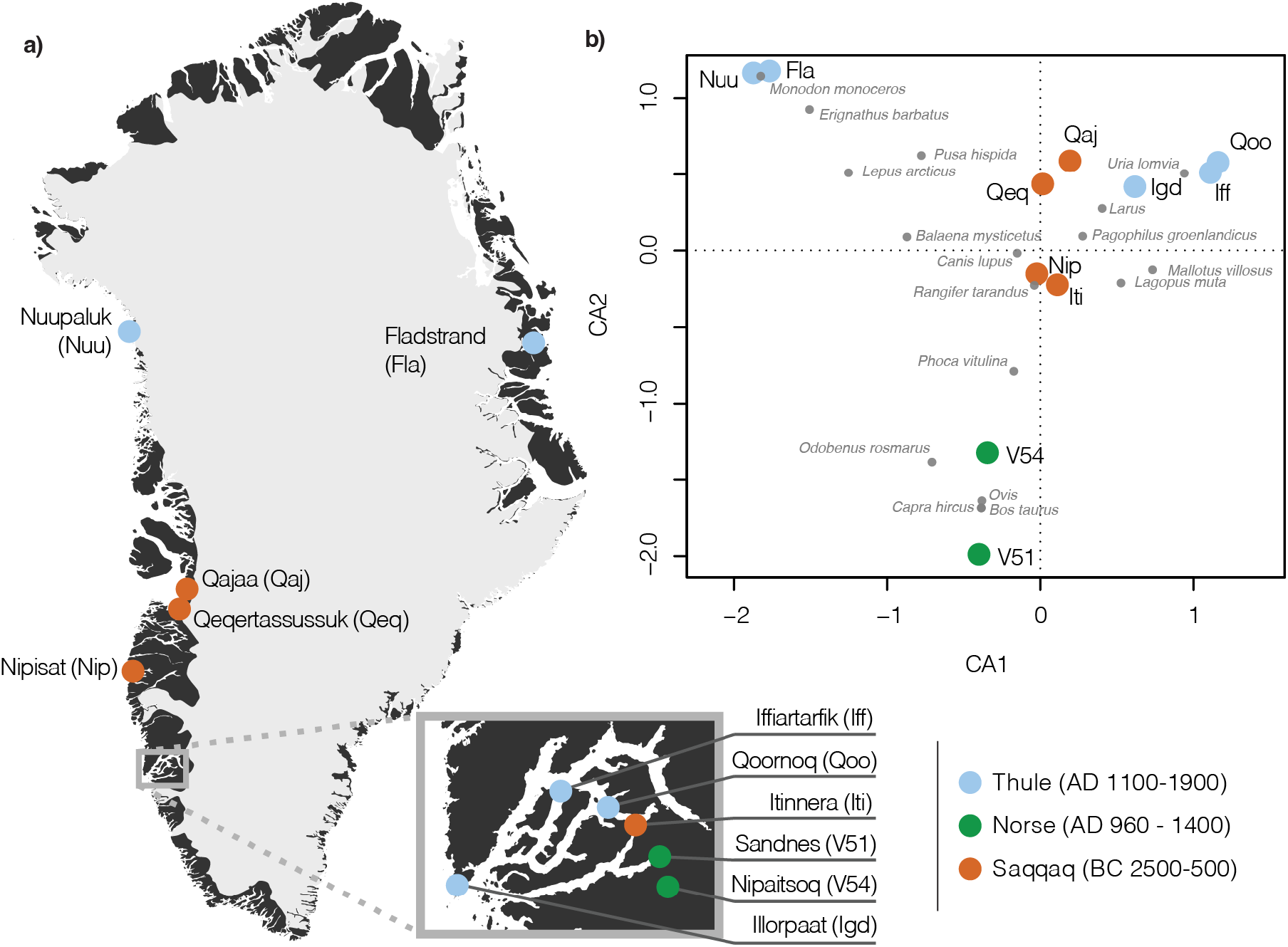
Sample location and diversity. **a)** Location of bulk bone samples. Circle colour indicates the cultural context for each sample. **b)** Coordination analysis of taxa identified at each site with BBM. Sites are depicted with large coloured circles, while taxa are represented by grey dots. For simplicity, only taxa with clear geographical patterns are depicted.

**Figure 2.**
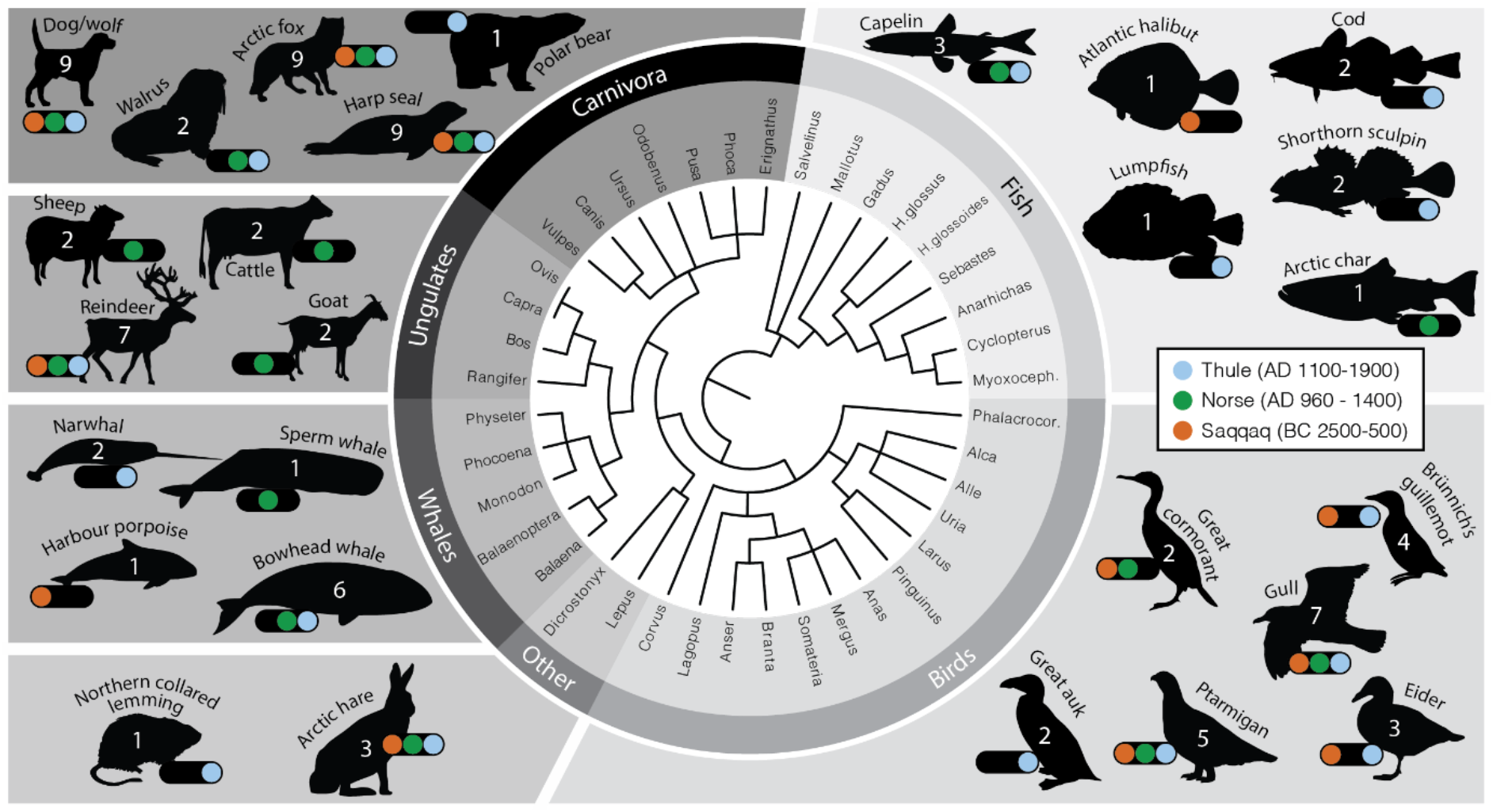
Species diversity detected from bulk bone samples. Dendrogram of genera detected in this study, with silhouettes of select species. Count in each silhouette represents the number of sites in which each taxon was detected (n=11). Narwhal, bowhead whale and sperm whale silhouette by <u>Chris Huh</u>.

Our data set is dominated by harp seal (*Pagophilus groenlandicus*; 10/11 sites), dog or wolf (*Canis lupus*; 9/11 sites) and arctic fox (*Vulpes lagopus*; 9/11 sites). The abundance of harp seal in our data is in agreement with morphological data where harp seal is typically detected in large quantities in middens across most of Greenland (with the exception of sites in Northern Greenland) ^2,20,21^. Conversely, morphological identification of dog remains, is fairly uncommon in Saqqaq middens but is more prominent in Norse, and, in particular, Thule culture sites. Lastly, although arctic fox is widespread in Greenland, fox remains are not normally identified zoo-archaeologically in abundances comparable to harp seal. The ubiquity of fox DNA across sites in our data could suggest that fox was an important resource to past cultures of Greenland. However, for both foxes and dogs, it is also possible that their scavenging on the midden waste could have left a genetic imprint in the form of urine or feces.

Generally, the species composition at each site appears to reflect both the geographical location of the site and the specific cultural practices of the people accumulating the bones (Figure 1). The two most distinct clusters in the coordination analyses are formed by (1) the two northern Thule sites, Fladstrand and Nuupaluk, and (2) the two Norse sites, Sandnes (V51) and Nipaatsoq (V54). The clustering of the northern sites is driven by the northern geographical ranges of species such as bearded seal (*Erignathus barbatus*) and narwhal (*Monodon monoceros*) which are found almost exclusively in northern sites. Furthermore, other species, such as the harbour seal (*Phoca vitulina*), are restricted to lower latitudes, and clusters within the southern sites (Figure 1). The clustering of the two Norse sites reflects their distinct subsistence strategy as farmers, and, accordingly, sheep, goat and cattle remains are abundant in the Norse middens^22^.

To validate our results, we also compared the metabarcoding approach with an alternative genetic approach based on shotgun metagenomics^16^. We sequenced shotgun libraries from 5 samples at two sites (Iffiartarfik; 2 samples and Qoornoq; 3 samples) and compared this data with metabarcoding data from the sites (Supplementary Figure 1; Supplementary Note 1). This comparison clearly highlights some of the commonly discussed pros and cons for both approaches^16,23^. We found that, despite sequencing 15% the number of reads (metabarcoding: 11.5 million reads, shotgun metagenomics: 74.2 million reads), metabarcoding detected a far greater species diversity (21 species) than shotgun metagenomics (11 species; Supplementary Figure 1a). Furthermore, all but one of the species (the great auk; *Pinguinus impennis*) detected by shotgun metagenomics was also detected by metabarcoding.

### Genetic evidence of past fishing practices in Greenland

The importance of fishing in ancient Greenland is generally understudied because of the paucity of fish bones in Greenlandic middens^2,24^. This has sparked a debate on the contribution of fishing to the overall diet for both the Paleo-Inuit^2^, Thule^25^ and Norse^24^. One of the advantages of the BBM approach is the ability to detect DNA from small and fragile bones that cannot be identified morphologically^26,27^. In our data, this is exemplified by the detection of nine different fish species, of which, four species (Atlantic wolffish; *Anarhichas lupus*, lumpfish; *Cyclopterus lumpus*, redfish; *Sebastes*, American plaice; *Hippoglossoides platessoides*) were not detected morphologically at any of the sites analysed. The two Thule sites in the Nuuk area Qoornoq and Iffiartarfik are particularly abundant in fish species, with 6 species detected at each site.

The diversity of fish species at these two sites might reflect the use of 4 mm mesh sieves during the excavation process, which is known to increase the number of fish bones excavated^28^. However, a combination of excellent preservation conditions and young age, might also have contributed to the high fish diversity at these sites.

The most common fish species identified in our data is the capelin (*Mallotus villosus*; 3/11 sites). As the capelin was most likely caught during the spawning season, their bone remains would have been tiny, which explains their absence from the morphological record. Capelin remains were identified at two Thule sites (Iffiartarfik and Qoornoq) and, for the first time, at the Norse site Sandnes. While capelin is a small fish it was likely an important economic element in both the Thule culture and for the Norse. They are arrive in such numbers that they can be scooped out of the water in large quantities, arriving at the same spawning grounds year after year^2^ and at a time (May) when winter supplies would be running low. Still, it should be kept in mind that some of the smaller fish species identified here, such as the capelin, may possibly be derived from stomach content of birds or seals brought to the site.

The diversity of the fish assemblage identified with BBM suggests the utilisation of a wide range of different fishing technologies. The presence of cod at two Thule sites in the Nuuk fjord indicates the use of lure or baited hooks. At the Norse site V54, we also find evidence of arctic char which spawns in rivers or lakes during the summer months. Situated by the Eqalunnguit (literally, ‘char river’), Arctic char was most likely caught by leister at this particular locality. This important resource is easily underestimated if the past economy is reconstructed from bone counts alone. Furthermore, end prongs from fishing leisters have also been identified from both Saqqaq^29^ and Thule cultural sites^25^, suggesting that leister fishing in shallow water along rivers, from the ice edge or from kayak was also a common Inuit practice in Greenland. Lastly, at the two fjord sites Iffiartarfik and Qoornoq, it is likely that net fishing contributed to the high number of fish species detected at both sites. Direct evidence of net fishing is not common in the archaeological record of Greenland, but net remains have been identified at a number of sites^29,30^ together with net sinkers^31–33^, and needles for net making^32^.

### Gulls, alcids, ducks and the extinct great auk

In the bird assemblage, we find evidence of utilization of a broad range of both gulls, alcids, duck and rock ptarmigan. The most commonly detected bird taxon is gulls (*Larus* sp; 7 sites), followed by sea ducks (*Somateria/Mergus* sp.; 6 sites) and rock ptarmigan (*Lagopus* muta; 5 sites). We also detect DNA from Brünnich’s guillemot (*Uria lomvia*) at the three Thule sites (Illorpaat, Iffiartarfik and Qoornoq) in Nuuk Fjord and one Paleo-Inuit site (Qajaa) in the Disko Bay. Detection of the genus *Branta* at the Nipisat site confirms previous morphometric studies of the Nipisat goose assemblage and supports the notion that during the Saqqaq period breeding distribution of Greenlandic geese differed from the present day ranges^34^. Lastly, from metagenomic data, we detect DNA from the extinct great auk (*Pinguinus impennis*) at two Thule culture sites from the Nuuk area. As demonstrated by Thomas et al. (2019)^35^, the extinction of the great auk was most likely caused by aggressive exploitation by European sailors. In the zoo-archaeological record, great auk remains associated with the Thule culture have exclusively been identified from coastal sites. Accordingly, it has been suggested that great auks were wintering off the coast of West Greenland between 1350 and 1800 AD^36^. However, we detect great auk DNA from the sites Qoornoq and Iffiartarfik which are both located deep in the Nuuk fjord system over 50 km from the coast. This finding suggests that great auks either made their way into the Nuuk fjord system, or that Thule bird hunters took the long journey to hunt this species. Furthermore, during the Saqqaq period, the presence of great auk was also documented deep in the Nuuk fjord at the Itinnera site where six great auk bones have been identified morphologically^36^.

### Detecting the invisible whale

We detected a total of five different whale species from seven sites (63% of sites analysed). The most commonly detected species was the bowhead whale (*Balaena mysticetus*; 6 sites), followed by the narwhal (*Monodon monoceros*; 2 sites). In addition to the bowhead whale, we detected two large whales: sperm whale (*Physeter catodon*) and fin whale (*Balaenoptera physalus*). The abundance of bowhead whale, and the detection of an additional two species of the largest whales is surprising because of the skill and effort required to hunt these taxa. It is possible that the detection of some of these species could be attributed to the scavenging of beached whale carcasses^16,32^. However, the consistent detection of bowhead whale across the majority of sites analysed suggests otherwise. When data from Seersholm et al. (2016) is included where bowhead whale DNA was detected from midden sediment at Qajaa and Qeqertassussuk as well, the total number of sites in this study with bowhead whale DNA adds up to 8 (73% of sites analysed). This indicates that bowhead whale was routinely exploited by all cultures in Greenland and could suggest the presence of a cooperative social structure aimed at big whale hunting that required the coordination of dozens of people to haul the animal ashore or onto the sea ice.

The presence of bowhead whale DNA deep in the Nuuk fjord at the two sites, Iffiartarfik and Qoornoq (dated to AD ~1500-1800), raises questions about the former range of this species. The modern range of bowhead whales does not extend as far south as the Nuuk area^41^, but records from the mid 1800’s suggest that it did sporadically in the past^42^. However, it is also possible that the presence of bowhead whale DNA at Iffiartarfik and Qoornoq is the result of trading, either with other Inuit or with European whalers who started operating in the area in larger numbers in the early 18^th^ century^43^.

To assess how the bowhead whale DNA detected here relates to the global bowhead whale population, we amplified short control region (CR) sequences with bowhead whale specific primers from the sites Nuupaluk, Iffiartarfik and Qoornoq (Supplementary Material, Table 3). By comparing the CR amplicon sequences with previously published data on 367 bowhead whale samples^37–40^, we found that the bowhead whale diversity in our samples reflects the structure in the general population. We identified the two most common bowhead whale CR haplotypes, at comparable frequencies as those observed in the general population (Figure 3). These data suggest that the population of bowhead whale hunted in Greenland both during Paleo-Inuit times and during the Thule culture does not differ genetically from today’s population. Furthermore, this finding confirms results from Borge et al. (2007)^37^ who found no clear geographical pattern of genetic diversity in bowhead whales when comparing the Svalbard stock with the Bering-Chukchi-Beaufort Seas stock. Our bulk bone aDNA findings also agree with data from Foote et al. (2013) who demonstrated genetic continuity in the bowhead whale lineages spanning the Pleistocene to Holocene^40^.

**Figure 3.**
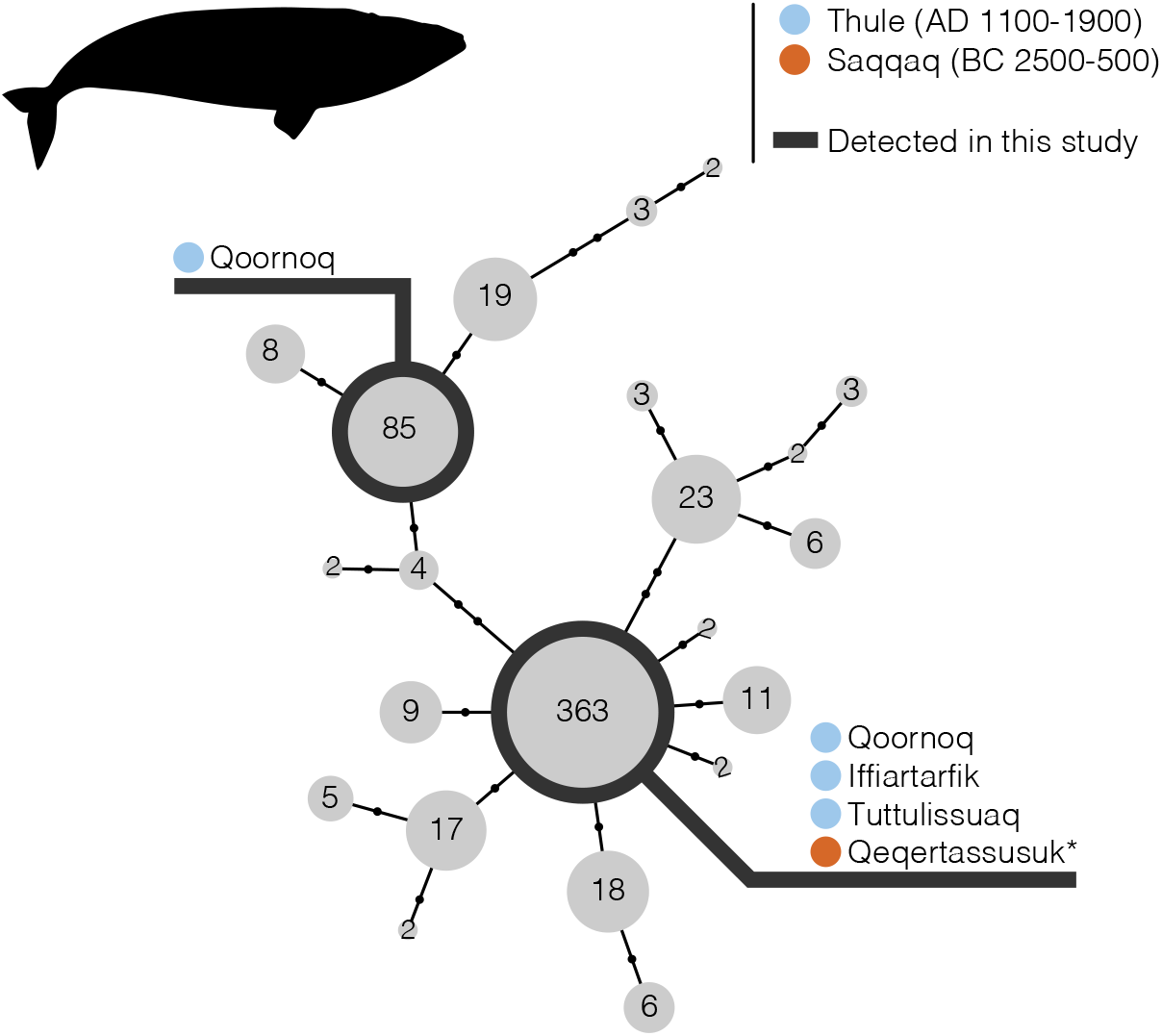
Bowhead whale diversity. Haplotype network based on 79 bp of the control region. Data compiled from Borge et al. (2007)^37^, Leduc et al. (2008)^38^, McLeod et al. (2012)^39^ and Foote et al. (2013)^40^, excluding singleton haplotypes. *From Seersholm et al. 2016^16^.

### A genetically distinct subgroup of caribou from the Saqqaq site of Itinnera

It has previously been suggested that a population of small caribou adapted to a high arctic environment were the first caribou to populate Greenland^44^. These “polar caribou” were probably descendants of the Peary caribou (*Rangifer tarandus pearyi*) and immigrated into Greenland from Ellesmere Island when the ice first retreated 9,000 years ago. At around 4,000 BP the larger tundra adapted subspecies of caribou (*Rangifer tarandus groenlandicus*) arrived in Greenland. During the next 2,000 years the tundra caribou spread across Greenland, and eventually outcompeted the polar caribou^45^.

Faunal remains of the polar adapted caribou are rare in Greenland, with the exception of Itinnera. This site was a Saqqaq caribou hunting camp, containing thousands of caribou bones from a subspecies significantly smaller than the modern population. In Meldgaard (1986) morphometric analyses of the Itinnera caribou revealed anomalies in the dentition on the jaws and relatively shortened legs, compared to the modern caribou population in Greenland. Based on these observations Meldgaard suggested that these animals belonged to a caribou population genetically divergent from the modern population. However, this hypothesis has never been corroborated by genetics.

With the high prevalence of caribou in our data (detected in 7 out of 11 sites and 29 out of 50 subsamples), we have a dataset of *12S* and *16S* caribou sequences with a reasonable temporal and spatial distribution, including the genetic data of the Itinnera caribou. We found that all four subsamples at Itinnera had the same novel *16S* haplotype, distinct from all other samples (Figure 4).

**Figure 4.**
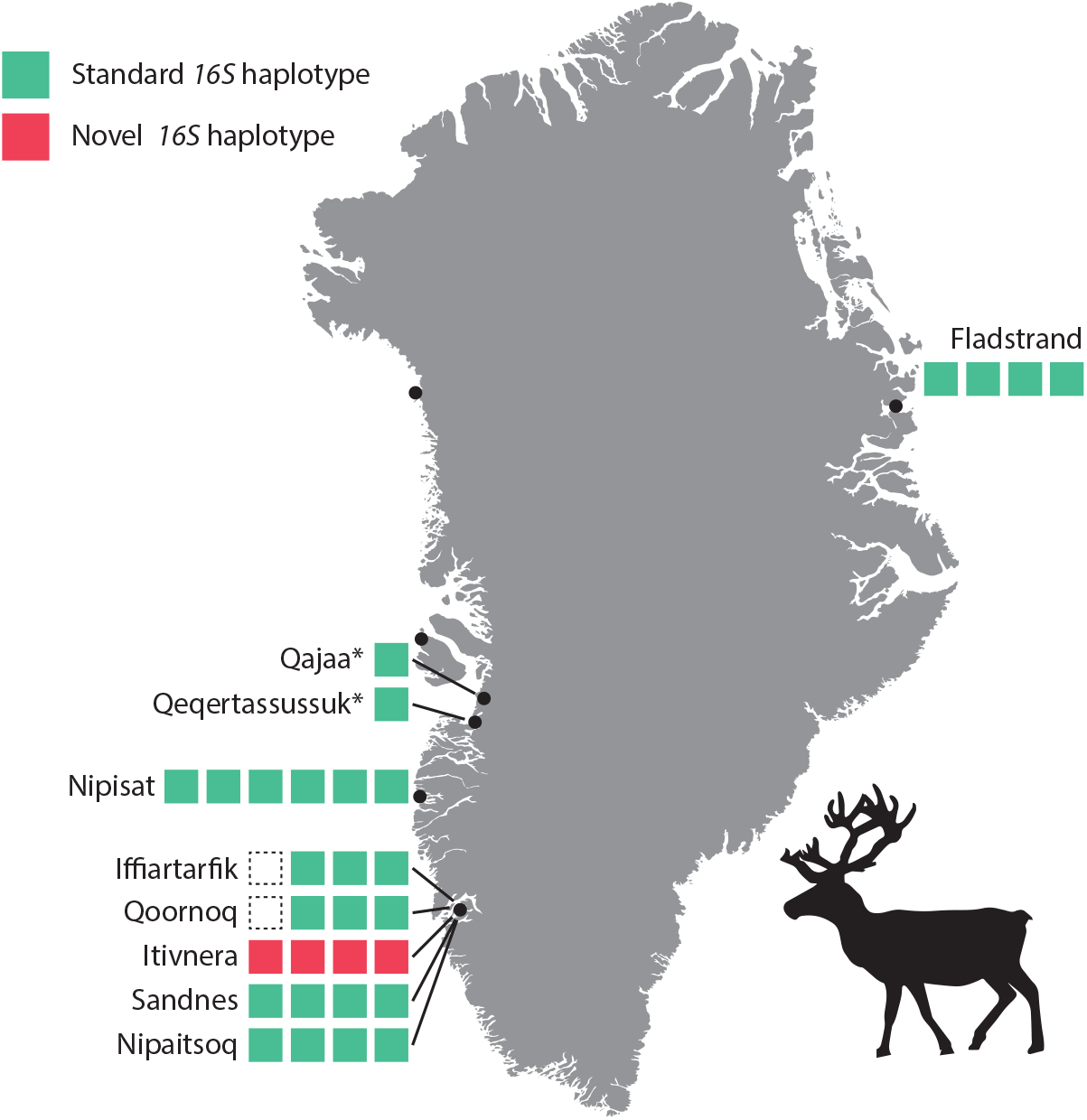
A novel 16S caribou haplotype from Itinnera. Variation in caribou 16S haplotypes across the sites analysed. Each coloured square represents one subsample of 50 bone fragments. Empty squares represent bulk bone samples in which caribou were not identified with the *16S* assay. *Haplotype identified from shotgun data from Seersholm et al. 2016^16^.

This finding strongly supports the hypothesis proposed by Meldgaard (1986) of an extinct, early settling, population of caribou present at Itinnera 4,000 years ago. Importantly, the novel Itinnera haplotype was not detected in the three other Saqqaq sites analysed (Nipisat, Qajaa, Qeqertassusuk), despite the fact that the sites are contemporaneous in age. This conforms with previous morphological analysis suggesting that the caribou from Nipisat, Qajaa and Qeqertassusuk were all comparable in size to the modern population^32,44^. Hence, during the Saqqaq period, the tundra caribou had colonised Western Greenland while the polar adapted caribou was still present in the Nuuk area further south. As previously suggested, it is likely that the Maniitsoq Ice Cap acted as a physical barrier for the spread of the tundra caribou southward. At some point over the following 2,000 years, the tundra caribou must have succeeded in crossing the ice cap, and subsequently outcompeted its polar adapted counterpart.

## Conclusion

Fish and whale remains are routinely assumed to be underrepresented in zoo-archaeological assemblages because of taphonomic processes. Our findings provide a rare window into whaling and fishing practices in ancient Greenland using genetics. In agreement with previous analyses from Greenland^16^ and Canada^46^, the most common whale species across the sites analysed is the bowhead whale. The preference for bowhead whales over other large whales is most likely explained by their low agility in combination with their relative high buoyancy, which ensured that the animal would remain afloat after being harpooned^47^.

The fish assemblage identified by bulk-bone metabarcoding adds five fish species to the list of previously identified taxa at the sites analysed and provide the first fish identification at the Norse site of Sandnes. Furthermore, we identify capelin as the most common fish in the assemblage, which due to its small size, is likely to be missed with traditional zoo-archaeological approaches, due to excavation and preservation biases. While the number of study sites with fish DNA identified in the present study still remains small, we have demonstrated the future potential of analysing unidentifiable fish remains from already excavated archaeological assemblages.

Lastly, we demonstrate the advantage of applying a genetics approach to study ancient bone assemblages by identifying a hitherto unknown genetic variant in the caribou population from the Saqqaq site of Itinnera. This finding confirms the hypothesis proposed by Meldgaard in 1986^44^, of a distinct population of caribou present 4,000 years ago in southwest Greenland. As a possible early settler in Greenland, this population possessed a smaller body size with a lower energy requirement, allowing them to survive in the barren landscape exposed after the ice retreated.

Currently, climate change is causing an extensive alteration of the Arctic environment with critical impacts on archaeological sites and buried remains^48^. Rising air temperatures, permafrost thaw and increased microbial degradation is considered one of the largest threats to the continued preservation of organic archaeological deposits^49^ including archaeological bones^50^. As a consequence, we are looking into a future where the abundance of well-preserved archaeological bones may decrease, making traditional zoo-archaeological approaches less feasible. In this study we have shown that bulk bone metabarcoding may serve as an alternate method that can be applied when bone fragments are small and unidentifiable. However, our method still requires well-preserved DNA, and currently very little is known about the degradation of DNA in the buried environment.

Collectively, these findings underpin the value of applying bulk bone metabarcoding broadly on bone assemblages across the Arctic and overlaying aDNA methods with traditional zoo-archaeological methods based on morphology. By identifying both whale and fish diversity in ancient Greenland, we provide a window into an understudied area of subsistence practices across Greenland. With the excellent preservation conditions in the Arctic, future biomolecular studies incorporating paleogenomics, ancient environmental DNA and bulk-bone metabarcoding have the potential to provide a more comprehensive picture of the spatial, temporal and cultural diversity in whaling and fishing practices observed across the Arctic.

## Methods

### Sample collection, extraction, amplification and sequencing

Samples were collected from the Natural History Museum of Denmark in 2019. Each sample represents two subsamples of 50 bone fragments, collected from the same stratigraphic context. For all studied sites except Itivsaalik at least two samples (2×2×50 bone fragments) were collected. To minimise contamination, samples were only handled while wearing gloves. However, none of the samples were excavated for the specific purpose of ancient DNA analysis, and accordingly, we expect some modern contamination, especially from the excavators.

All laboratory processing of the samples was conducted at Curtin University, Western Australia. Upon import to Australia, samples were transported directly to the quarantine approved ultra clean laboratory facility TRACE (Trace Research Advanced Clean Environment) at Curtin University. Each subsample was ground to a fine bone powder on a PM200 planetary ball mill (Retsch) and split in two samples of 100-150 mg bone powder for extraction (extraction A and B). For each extraction bone powder was digested over night at 55°C under rotation in 1mL digestion buffer (0.25 mg/mL Proteinase K in 0.5M EDTA). Next, samples were centrifuged and the supernatant transferred to a MWCO 30 kDa Vivaspin 500 column (Merck) on which the supernatant was concentrated to 50-100 μL. Lastly, the concentrate was cleaned on a MinElute spin column (Qiagen) using a modified binding buffer consisting of 40% Isopropanol, 0.05% Tween 20, 90 mM NaAc and 5 M Guanidine hydrochloride^51^, but otherwise following the manufacturer’s instructions.

PCR amplifications were carried out in the following reaction concentrations: 1uL DNA in 1 x Gold PCR buffer, 1 mM MgCl_2_, 0.25 μM dNTPs, 2.5 U AmpliTaq Gold (Applied Biosystems), 0.12 X SYBR green (Invitrogen) and 0.4 mg/mL Bovine Serum Albumin (Fisher). All extracts were amplified with all four assays (Supplementary Material, Table 3) on a quantitative StepOnePlus PCR thermocycler (Thermo Scientific) with the following cycling conditions: 5 minutes at 95°C followed by 50 cycles of: 95°C for 30 seconds, 54-57°C (Supplementary Material, Table 3) for 30 seconds and 72°C for 1 minute, followed by a final elongation step of 72°C for 10 minutes. After amplification, samples were pooled by Ct-value in pools of approximately 16 reactions. Next, pools were blended at equimolar amounts based on DNA concentration readings from a QIAxcel capillary electrophoresis device (Qiagen). Lastly, the DNA library was size selected to retain only reads between 160 bp and 450 bp on a Pippin Prep (Sage Science). The final library was sequenced on the Illumina MiSeq platform with single end chemistry for 325 cycles using a standard flow cell (V3 chemistry).

### Bioinformatics

Reads were assigned to samples based on both tag and amplification primer sequence using the ngsfilter program in Obitools^52^. For each sample, reads were collapsed into unique reads (obiuniq) and reads shorter than 80 bp or represented by fewer than 5 reads in a sample were discarded (obigrep). Each sample was filtered to remove PCR artefacts using obiclean (obiclean -r 0.2 -d 2 -H), followed by two steps of sumaclust: “sumaclust -R 0.5 -t 0.95” and “sumaclust -R 0.01 -t 0.93”. Lastly, chimeric sequences were remoced with uchime-denovo from vsearch^53^. These steps greatly reduce the number of spurious reads identified in each sample, and with a maximum of 19 ASVs identified in a single sample of 50 bone fragments (Supplementary Material, Table 4 and Table 5), we are confident that the filtering is sufficient to reveal the true biodiversity in our samples.

Species assignments were carried out on the filtered ASVs by querying each sequence against the NCBI nt database using megablast^54^. Blast files were parsed using the python script <u>blast_getLCA</u>^55^ which assigns each read to the lowest common ancestor of the best hit(s) in the database. Lastly, raw species assignments were parsed manually, and each assignment was correlated with known species occurrences in Greenland. For example, if a sequence had equal identity to two species of which only one is known to occur in Greenland, only the species present in Greenland would be considered for the taxonomic assignment.

### Statistics and data visualisation

Correspondence analysis (Figure 1b) was carried out on presence/absence data of identified taxa (Supplementary Material, Table 8) using the R package vegan (https://cran.r-project.org/web/packages/vegan/). The bowhead whale diversity analysis (Figure 3) was based on data from Borge et al. (2007)^37^, Leduc et al. (2008)^38^, McLeod et al. (2012)^39^, Foote et al. (2013)^40^, and Seersholm et al. 2016^16^ excluding singleton haplotypes. The haplotype network was constructed using the function haploNet (pegas) in R.

### Shotgun sequencing

DNA sequencing libraries were built using a single stranded approach^56^ as described in Grealy et al. (2017)^57^, and sequenced on the Illumina NextSeq system at Curtin University. Bioinformatics were conducted following the pipeline described in Seersholm et. al (2016). In brief, adapter sequences were trimmed and paired-end reads were merged using AdapterRemoval (v. 2.3.0)^58^, discarding sequences shorter than than 25 bp. Next, low complexity reads with a dust threshold higher than 1.0 were removed with sga preprocess. This step was implemented to ensure that low complexity DNA in the samples would not result in false positive assignments to species with high contents of simple repeats in the database. For taxonomic assignments, reads preprocessed by sga were mapped against the NCBI refseq database of full mitochondrial genomes (ftp://ftp.ncbi.nlm.nih.gov/refseq/release/mitochondrion/) using bowtie2^59^ set to report up to 500 hits per sequence read. The resulting sam files were then parsed using the getLCA script (https://github.com/frederikseersholm/getLCA), which assigns each read to the taxonomic node of the lowest common ancestor(s) of the best hits to the database.

## End Notes

## Acknowledgements

This study was supported by the Australian Research Council Discovery Project DP160104473 and Forrest Research Foundation (to F.V.S.). Data analysis was carried out with support from the Pawsey Supercomputing Centre. Furthermore, the authors would like to thank Henning Matthiesen and the “REMAINS of Greenland” project funded by the VELUX FOUNDATION, and Kiristian Gregersen and the Natural History Museum of Denmark.

## Author contributions

F.V.S., M.B., A.J.H. and M.M. designed the study. F.V.S. collected samples, conducted laboratory work, analysed the data and wrote the manuscript with input from all co-authors.

