## Supplemental Material for "New insights into 4,000 years of resource economy across Greenland using ancient DNA"

### Supplementary Figures

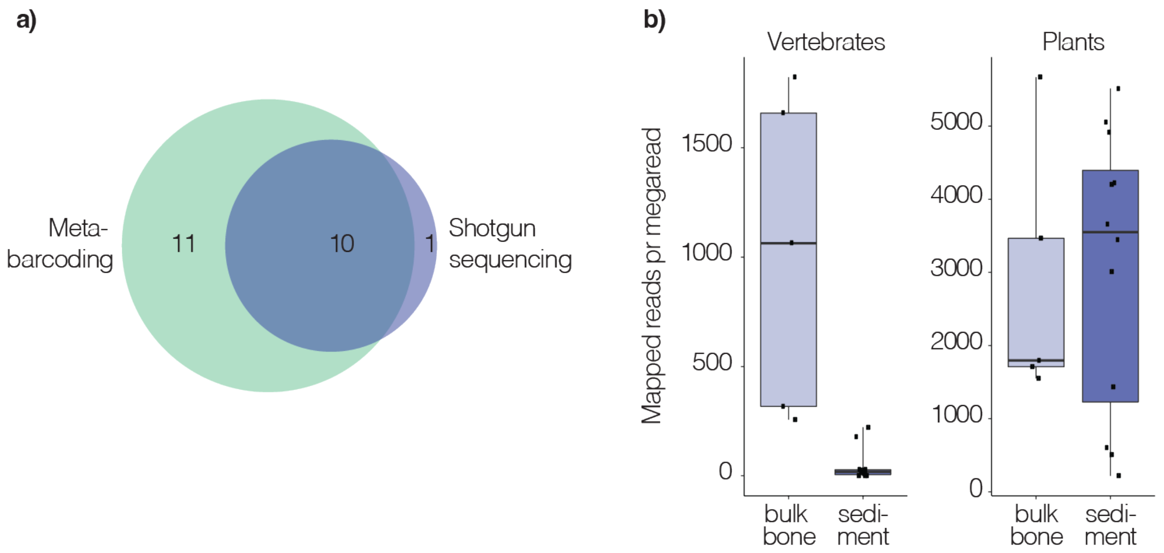


**Supplementary Figure 1. Comparison between BBM and shotgun sequencing.** a) Venn diagram comparing the number of species detected by bulk bone metabarcoding (BBM) and by shotgun sequencing. b) A comparison of the abundance estimates from metabarcoding of the three most abundant species detected in the 12Sv5 assay with their abundances as estimated by shotgun metagenomics. Metabarcoding results are presented as green boxplots since each bulk bone sample was analysed with metabarocding in replicates of 8. Shotgun sequencing results are presented as blue diamonds. c) Boxplots comparing the number of assigned reads per megaread sequenced for vertebrates and plants, respectively, using bulk bone (this study) or sediment (Seersholm et. al 2016[1]) as substrate.

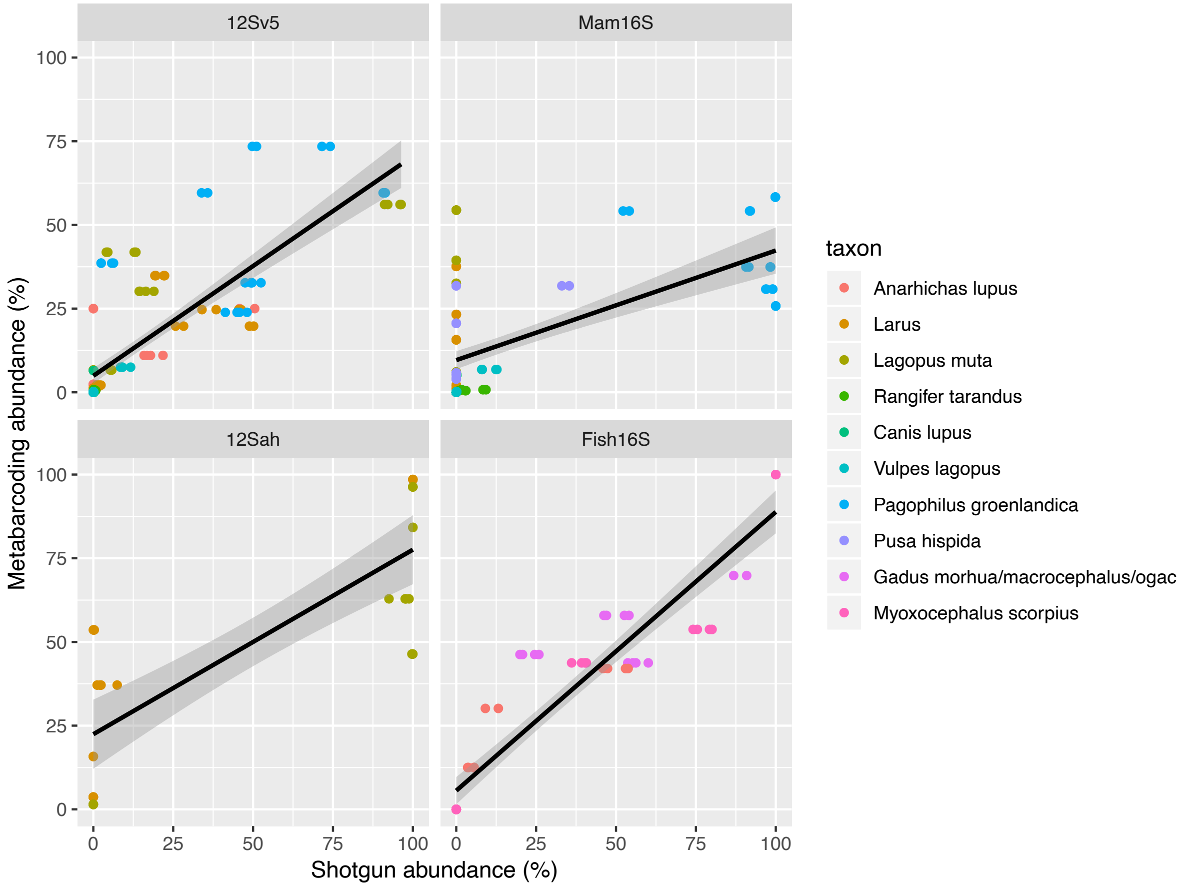


**Supplementary Figure 2. Abundance relationship of taxa detected by both metabarcoding and shotgun sequencing.** For each assay the relative abundance was calculated for species detected by both approaches. Data is plotted for each assay.

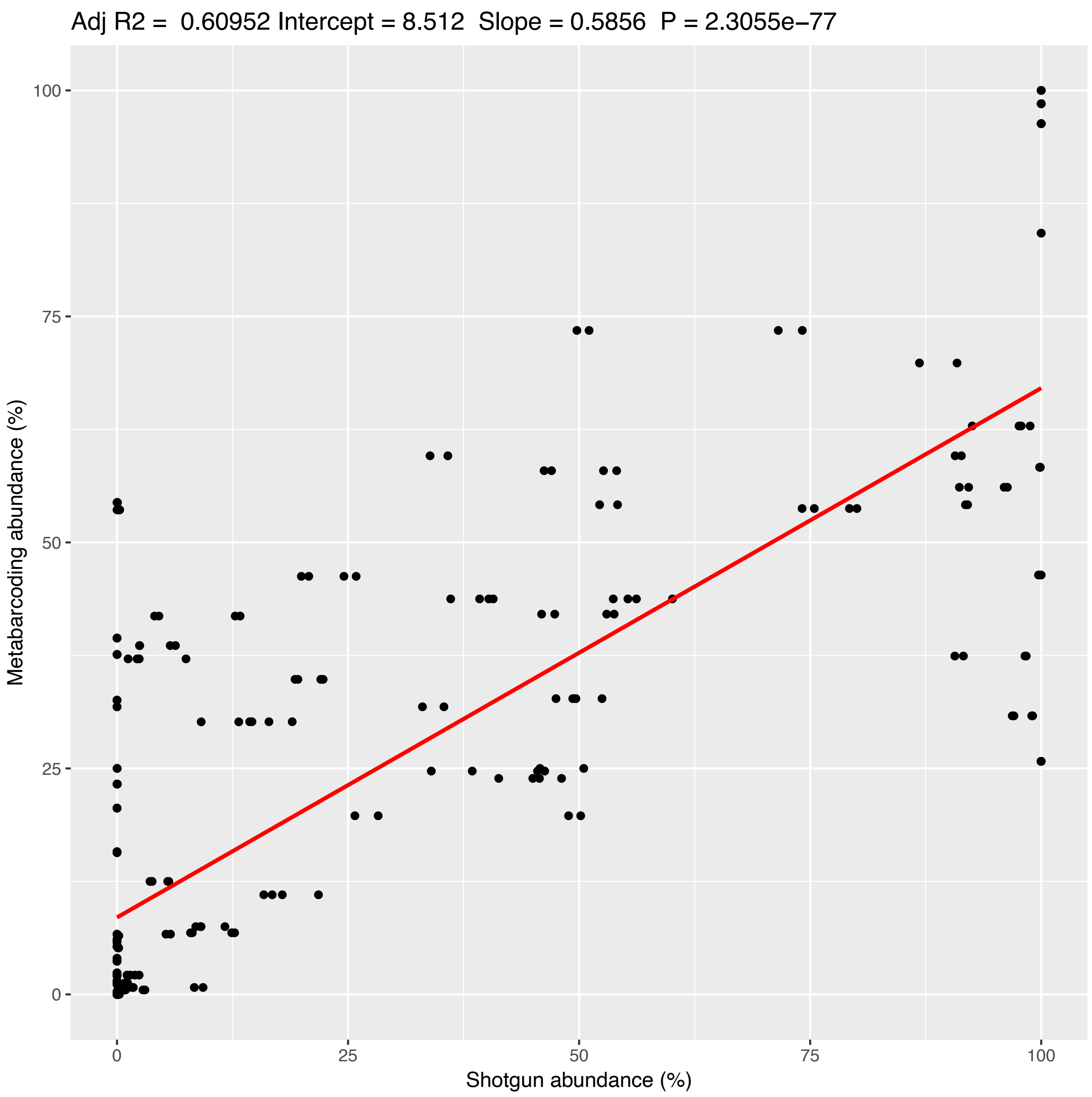


**Supplementary Figure 3. Abundance relationship of taxa detected by both metabarcoding and shotgun sequencing.** For each assay the relative abundance was calculated for species detected by both approaches. Data from each assay (Supplementary Figure 2) was merged to a single plot.

Supplementary Tables

**Table 1. Sample information.** *No endogenous DNA detected.

| ID | Site | Location/context | n bones |
|---|---|---|---|
| Qoo1 | Qoornoq | x230, 40-50 cm | 100 |
| Qoo2 | Qoornoq | x225, 30-40 cm | 100 |
| Qoo3 | Qoornoq | x249, 60-70 cm | 100 |
| Iff1 | Iffiartarfik | x502, 0-20 cm | 100 |
| Iff2 | Iffiartarfik | x536, 20-30 cm | 100 |
| Nip1 | Nipisat | 384/188, lag 2k (phase 1) | 100 |
| Nip2 | Nipisat | 383/192, lag 2k (phase2) | 100 |
| Nip3 | Nipisat | 398/205, lag 2k (phase3) | 100 |
| Igd1 | Illorpaat | Felt D, lag 15, Plan XVI. V257, Ø262 | 100 |
| Igd2 | Illorpaat | Felt C, plan 2 Mødding. | 100 |
| Nuu1 | Nuupaluk | F3 Lag 9 | 100 |
| Nuu2 | Nuupaluk | F3 Lag 7 | 100 |
| Iti1 | Itinnera | Gruppe A | 100 |
| Iti2 | Itinnera | Gruppe B | 100 |
| Itv1* | Itivsaalik | Hus II | 100 |
| V541 | Nipaatsoq | 492/194, 320-325 ØR. Ni V54 76-2 | 100 |
| V542 | Nipaatsoq | 497/187, 360-375 | 100 |
| Qaj1 | Qajaa | Felt E,b 90-100 | 100 |
| Qaj2 | Qajaa | Felt E 70-80 | 100 |
| V511 | Sandnes V51 | P8IV, Su43, L7m, m7m? scrap | 100 |
| V512 | Sandnes V51 | 1091, P8 su 50 I, scrap, L7M, MTO? | 100 |
| Qeq1 | Qeqertassussuk | Qt 86 Felt B 13/23:54 | 100 |
| Qeq2 | Qeqertassussuk | Qt 90 Felt B 26/21.5:3 | 100 |
| Fla1 | Fladstrand (Cla-06) | 99,5/202,0 lag 2, unidentifiable | 100 |
| Fla2 | Fladstrand (Cla-06) | 98,5/200,0 lag 1, unidentifiable | 100 |

**Table 2. Archaeological sites.** *No endogenous DNA detected. ~ Indicate age estimates that are not based on radiocarbon dates, but on associated artefacts.

| ID | Site | Culture | Age | Latitude | Longitude | ZMK ID | NKAH ID | Reference |
|---|---|---|---|---|---|---|---|---|
| Nuu | Nuupaluk | Neo-Inuit (Historic Inuit) | AD 1600 - 1800 | 75.341709 | -58.649506 | 81/1979 | NKAH 3737 | Meldgaard & Grønnow 1980[2] |
| Fla | Fladstrand | Neo-Inuit (Thule) | AD 1400 - 1850 | 74.098703 | -21.187391 | 101/2007 | NKAH 4300 | Gotfredsen 2010[3] |
| Igd | Illorpaat | Neo-Inuit (Historic Inuit) | AD 1500 - 1800 | 64.12388 | -52.08371 | 142/1972 | NKAH 1081 | Gulløv 1997[4], Møhl 1980[5] |
| Qaj | Qajaa | Paleo-Inuit (Saqqaq) | 1900 - 900 BC | 69.127700 | -50.705796 | 350/1982 | NKAH 1794 | Møhl 1986[6] |
| Qeq | Qeqertassussuk | Paleo-Inuit (Saqqaq) | 2400-1400 BC | 68.593000 | -51.072395 | 70/1983 | NKAH 1613 | Meldgaard 2004[7] |
| Nip | Nipisat | Paleo-Inuit (Saqqaq) | 1890 - 990 BC | 66.81457 | -53.492956 | 136/1989 | NKAH 307 | Gotfredsen & Møbjerg 2004[8] |
| Itv* | Itivsaalik | Neo-Inuit (Thule) | AD 1500 - 1650 | 64.11327 | -52.08758 | 143/1972 | NKAH 1085 | Gulløv 1997[4] |
| Qoo | Qoornoq | Neo-Inuit (Historic Inuit) | ~AD 1700 - 1800 | 64.533710 | -51.086536 | - | NKAH 577 | Hans Harmsen, unpublished |
| Iff | Iffiartarfik | Neo-Inuit (Historic Inuit) | ~AD 1500 - 1700 | 64.461740 | -50.649667 | - | NKAH 1202 | Hans Harmsen, unpublished |
| Iti | Itinnera | Paleo-Inuit (Saqqaq) | 1000 BC | 64.383333 | -50.399997 | 114/1966, 112/1958 | NKAH 1199 | Møhl 1972[9] |
| V51 | Sandnes (V51) | Norse | AD 1050 - 1350 | 64.242640 | -50.182338 | 75/1981 | NKAH 1480 | McGovern et al. 1996[10] |
| V54 | Nipaatsoq (V54) | Norse | AD 1030 - 1440 | 64.107710 | -50.112158 | 137/1977 | NKAH 1486 | McGovern et al. 1983[11] |

**Table 3. Primers**

| | Forward Primer | Reverse Primer | Annealing Temp | ref |
|---|---|---|---|---|
| 12SV5 | TAGAACAGGCTCCTCTAG | TTAGATACCCCACTATGC | 57°C | Riaz et al. (2011)[12] |
| Mam16S | CGGTTGGGGTGACCTCGGA | GCTGTTATCCCTAGGGTAACT | 57°C | Taylor (1996)[13] |
| Fish16S | GACCCTATGGAGCTTTAGAC | CGCTGTTATCCCTADRGTAACT | 54°C | Deagle et al. (2007)[14], Berry et al. (2017)[15] |
| 12SAH | CTGGGATTAGATACCCCACTAT | CCTTGACCTGTCTTGTTAGC | 57°C | Cooper (1994)[16] |
| BM_short | TTCACTACGGGAAGTTAAAGCTCG | GGAGCGGCCATAGGATTCAGTTG | 54°C | Foote et al. (2013)[17] |

### Table 4. Sequencing information, assays 12SV5 and Mam16S.

| Sample | 12SV5 | | | | | | Mam16S | | | | | |
|---|---|---|---|---|---|---|---|---|---|---|---|---|
| | Extraction 1 | | | Extraction 2 | | | Extraction 1 | | | Extraction 2 | | |
| | Raw count | Filt. count | Uniq. count | Raw count | Filt. count | Uniq. count | Raw count | Filt. count | Uniq. count | Raw count | Filt. count | Uniq. count |
| **Iff1_A_I** | 36276 | 26720 | 9 | 93490 | 71593 | 9 | 101805 | 84949 | 7 | 109616 | 73313 | 4 |
| **Iff1_A_II** | 21705 | 16049 | 8 | 103853 | 80152 | 9 | 114461 | 96191 | 5 | 122270 | 76394 | 5 |
| **Iff1_B_I** | 38388 | 30605 | 6 | 136996 | 106161 | 7 | 106487 | 87980 | 4 | 113604 | 90233 | 5 |
| **Iff1_B_II** | 43122 | 34412 | 7 | 128345 | 100029 | 9 | 125152 | 102740 | 5 | 129033 | 102963 | 4 |
| **Iff2_A_I** | 37274 | 26042 | 6 | 112556 | 83519 | 8 | 95317 | 73208 | 5 | 121576 | 99263 | 4 |
| **Iff2_A_II** | 78341 | 54653 | 7 | 126006 | 93339 | 8 | 120649 | 94202 | 6 | 117615 | 95569 | 4 |
| **Iff2_B_I** | 47195 | 34243 | 9 | 118604 | 84724 | 13 | 89277 | 52856 | 12 | 99533 | 55515 | 12 |
| **Iff2_B_II** | 36377 | 26371 | 7 | 108697 | 78234 | 11 | 91431 | 53248 | 13 | 94138 | 50689 | 10 |
| **Qoo1_A_I** | 28621 | 15642 | 6 | 105154 | 72674 | 9 | 91052 | 61826 | 6 | 98949 | 73416 | 6 |
| **Qoo1_A_II** | 23371 | 13086 | 7 | 96508 | 67644 | 9 | 87835 | 60914 | 10 | 97847 | 70878 | 7 |
| **Qoo1_B_I** | 33202 | 24564 | 9 | 121803 | 73012 | 14 | 98715 | 67875 | 8 | 120358 | 78723 | 8 |
| **Qoo1_B_II** | 33275 | 22663 | 7 | 117318 | 69567 | 13 | 92647 | 68379 | 9 | 93759 | 61416 | 9 |
| **Qoo2_A_I** | 51287 | 33045 | 7 | 105501 | 74179 | 7 | 65403 | 46964 | 3 | 109807 | 77564 | 3 |
| **Qoo2_A_II** | 33115 | 21937 | 6 | 117743 | 81930 | 6 | 98151 | 68624 | 3 | 114155 | 81206 | 2 |
| **Qoo2_B_I** | 29875 | 22152 | 7 | 111570 | 77902 | 12 | 105724 | 93571 | 8 | 116010 | 96153 | 8 |
| **Qoo2_B_II** | 29103 | 21383 | 8 | 116810 | 82536 | 11 | 115813 | 102575 | 8 | 116058 | 96600 | 7 |
| **Qoo3_A_I** | 55034 | 34068 | 6 | 131944 | 79959 | 7 | 113553 | 93888 | 4 | 111551 | 93389 | 3 |
| **Qoo3_A_II** | 47534 | 29170 | 5 | 120152 | 72147 | 7 | 106261 | 89154 | 3 | 102824 | 86058 | 2 |
| **Qoo3_B_I** | 48606 | 37735 | 8 | 134256 | 102338 | 11 | 142344 | 124701 | 2 | 104605 | 89136 | 2 |
| **Qoo3_B_II** | 45365 | 34798 | 8 | 124687 | 94278 | 12 | 89045 | 78514 | 2 | 113683 | 96529 | 2 |
| **Fla1_A** | 65447 | 56091 | 4 | 100170 | 78441 | 5 | 69910 | 59930 | 6 | 66308 | 50300 | 8 |
| **Fla1_B** | 66763 | 34202 | 7 | 60430 | 23283 | 4 | 69221 | 31423 | 7 | 45648 | 21660 | 9 |
| **Fla2_A** | 46117 | 25291 | 12 | 63712 | 34036 | 10 | 63028 | 47908 | 13 | 52993 | 42132 | 12 |
| **Fla2_B** | 56245 | 39921 | 8 | 59152 | 41998 | 10 | 106394 | 87658 | 12 | 106723 | 89476 | 9 |
| **Igd1_A** | 41128 | 35864 | 1 | 84831 | 73944 | 1 | 62101 | 58019 | 3 | 55766 | 49993 | 3 |
| **Igd1_B** | 68629 | 56648 | 3 | 81014 | 61777 | 4 | 191878 | 174669 | 3 | 56421 | 51131 | 2 |
| **Igd2_A** | 51709 | 45522 | 1 | 41619 | 36936 | 1 | 145791 | 138173 | 1 | 67928 | 63972 | 1 |
| **Igd2_B** | 78467 | 68199 | 1 | 45324 | 39217 | 3 | 66149 | 62494 | 2 | 62723 | 58790 | 1 |
| **Iti1_A** | 52266 | 45185 | 4 | 94669 | 80638 | 5 | 73580 | 68339 | 4 | 64777 | 59387 | 4 |
| **Iti1_B** | 51257 | 25887 | 4 | 62098 | 52653 | 3 | 76601 | 71681 | 3 | 60202 | 46565 | 3 |
| **Iti2_A** | 91189 | 81179 | 2 | 14316 | 13081 | 3 | 112772 | 99204 | 2 | 64425 | 51575 | 5 |
| **Iti2_B** | 6313 | 5761 | 1 | - | - | - | 84042 | 78381 | 3 | 101115 | 63992 | 1 |
| **Itv1_A** | 47969 | 38190 | 2 | 39338 | 31929 | 2 | 91268 | 86998 | 1 | 62713 | 56048 | 2 |
| **Itv1_B** | - | - | - | 42125 | 37794 | 1 | 89008 | 85112 | 1 | 130132 | 118549 | 1 |
| **Nip1_A** | 36167 | 18395 | 9 | - | - | - | 93266 | 67311 | 5 | 56177 | 42192 | 4 |
| **Nip1_B** | 31871 | 16764 | 9 | 61466 | 42359 | 7 | 68926 | 53684 | 7 | 42189 | 33606 | 7 |
| **Nip2_A** | 42950 | 22486 | 5 | 51788 | 25756 | 6 | 68247 | 47689 | 3 | 68079 | 47741 | 5 |
| **Nip2_B** | 52028 | 34405 | 8 | - | - | - | 94440 | 64804 | 3 | - | - | - |
| **Nip3_A** | 43607 | 21295 | 4 | 45828 | 22482 | 7 | 82979 | 52095 | 4 | 101972 | 57506 | 5 |
| **Nip3_B** | 42498 | 25972 | 6 | 39443 | 24795 | 7 | 100344 | 69215 | 7 | 89809 | 63708 | 5 |
| **Nuu1_A** | 70661 | 45247 | 6 | 71347 | 33182 | 6 | 111135 | 73829 | 7 | 70221 | 43229 | 7 |
| **Nuu1_B** | 59483 | 41002 | 3 | 70748 | 46538 | 4 | 49439 | 43777 | 4 | 85500 | 78280 | 1 |
| **Nuu2_A** | 60110 | 48472 | 4 | 68899 | 59111 | 1 | 104538 | 93068 | 6 | 47523 | 42644 | 2 |
| **Nuu2_B** | 37245 | 31605 | 1 | 78884 | 68268 | 1 | 49574 | 46594 | 1 | 56541 | 51919 | 2 |
| **Qaj1_A** | 55520 | 44628 | 4 | 96576 | 68390 | 3 | 55554 | 43016 | 4 | 42584 | 26993 | 2 |
| **Qaj1_B** | 80214 | 59034 | 4 | 46764 | 31312 | 6 | 55333 | 36621 | 3 | 42694 | 32270 | 7 |
| **Qaj2_A** | 61057 | 54048 | 1 | 32705 | 28833 | 1 | 98419 | 93397 | 1 | 117406 | 105678 | 1 |
| **Qaj2_B** | - | - | - | 43571 | 37088 | 1 | 62212 | 59100 | 1 | - | - | - |
| **Qeq1_A** | 63621 | 32537 | 8 | 73499 | 42484 | 5 | 66171 | 52382 | 3 | 46175 | 32614 | 3 |
| **Qeq1_B** | 13752 | 4904 | 2 | 87487 | 73735 | 4 | 63199 | 27568 | 2 | 104986 | 91543 | 2 |
| **Qeq2_A** | 80052 | 57707 | 2 | 59951 | 47686 | 5 | 54967 | 24421 | 2 | 55832 | 33327 | 2 |
| **Qeq2_B** | 53030 | 43875 | 4 | 93088 | 52828 | 3 | 53765 | 24997 | 3 | 54540 | 33970 | 4 |
| **V511_A** | 51737 | 27446 | 10 | 57678 | 30592 | 8 | 50128 | 28692 | 7 | 87329 | 51773 | 10 |
| **V511_B** | 54747 | 35623 | 12 | 61512 | 40163 | 7 | 141974 | 107838 | 8 | 97563 | 75200 | 8 |
| **V512_A** | 64642 | 50982 | 7 | 71993 | 56610 | 6 | 156059 | 112248 | 13 | 47894 | 40244 | 7 |
| **V512_B** | 58265 | 44747 | 11 | 47403 | 31754 | 10 | 72002 | 56234 | 6 | 49275 | 36070 | 6 |
| **V541_A** | 36328 | 14869 | 9 | 57767 | 21814 | 10 | 57723 | 27157 | 11 | 60525 | 26346 | 13 |
| **V541_B** | 32934 | 13573 | 14 | 70505 | 31090 | 11 | 55518 | 26413 | 9 | 61647 | 31023 | 8 |
| **V542_A** | 40544 | 14515 | 10 | 70823 | 23912 | 9 | 54790 | 23199 | 10 | 48638 | 19664 | 11 |
| **V542_B** | 70014 | 47519 | 7 | 81463 | 55878 | 6 | 67519 | 52018 | 7 | 64737 | 51348 | 7 |

### Table 5. Sequencing information, assays Fish16S and 12SAH.

| Sample | Fish16S | | | | | | 12SAH | | | | | |
|---|---|---|---|---|---|---|---|---|---|---|---|---|
| | Extraction 1 | | | Extraction 2 | | | Extraction 1 | | | Extraction 2 | | |
| | Raw count | Filt. count | Uniq. count | Raw count | Filt. count | Uniq. count | Raw count | Filt. count | Uniq. count | Raw count | Filt. count | Uniq. count |
| **Iff1_A_I** | 39666 | 21859 | 4 | 109793 | 64383 | 4 | 64992 | 46099 | 1 | 124583 | 93611 | 4 |
| **Iff1_A_II** | 51701 | 26182 | 4 | 69499 | 42480 | 4 | 70561 | 49883 | 1 | 72750 | 53860 | 4 |
| **Iff1_B_I** | 38845 | 25279 | 4 | 72646 | 52855 | 4 | 58465 | 36509 | 6 | 36521 | 21398 | 2 |
| **Iff1_B_II** | 52463 | 34297 | 2 | 82646 | 58521 | 3 | 77754 | 48062 | 6 | 37855 | 24202 | 2 |
| **Iff2_A_I** | 77216 | 43137 | 2 | 82501 | 58282 | 2 | 75964 | 46271 | 5 | 73483 | 48650 | 3 |
| **Iff2_A_II** | 93783 | 49922 | 2 | 77446 | 54026 | 2 | 119370 | 67083 | 7 | 29256 | 10648 | 2 |
| **Iff2_B_I** | 31509 | 20696 | 2 | 70954 | 51358 | 2 | 77694 | 49673 | 5 | 45260 | 33500 | 2 |
| **Iff2_B_II** | 28870 | 17201 | 2 | 70430 | 53331 | 2 | 73635 | 49071 | 4 | 80025 | 57216 | 4 |
| **Qoo1_A_I** | 56357 | 34195 | 2 | - | - | - | 41377 | 31156 | 1 | - | - | - |
| **Qoo1_A_II** | 42062 | 25671 | 2 | - | - | - | 35136 | 24110 | 1 | 1 | - | - |
| **Qoo1_B_I** | 60110 | 37050 | 1 | - | - | - | 29502 | 16163 | 3 | - | - | - |
| **Qoo1_B_II** | 54418 | 30449 | 1 | 52 | 14 | 1 | 37835 | 14222 | 2 | - | - | - |
| **Qoo2_A_I** | 44074 | 27915 | 1 | 73724 | 60954 | 1 | 44829 | 15165 | 4 | 29860 | 16730 | 2 |
| **Qoo2_A_II** | 46822 | 30415 | 1 | 60531 | 49388 | 1 | 49103 | 15548 | 5 | 19322 | 11893 | 4 |
| **Qoo2_B_I** | 44090 | 20383 | 1 | 4 | - | - | - | - | - | - | - | - |
| **Qoo2_B_II** | 55501 | 21989 | 2 | 2 | - | - | - | - | - | - | - | - |
| **Qoo3_A_I** | 67917 | 35871 | 2 | 90156 | 55090 | 2 | 65653 | 41193 | 9 | 43782 | 30356 | 4 |
| **Qoo3_A_II** | 109305 | 54494 | 4 | 73405 | 45379 | 2 | 72220 | 44077 | 9 | 35933 | 27452 | 1 |
| **Qoo3_B_I** | 39097 | 21606 | 3 | 85427 | 53576 | 3 | 86395 | 41043 | 6 | 64662 | 35407 | 4 |
| **Qoo3_B_II** | 29741 | 16401 | 3 | 70035 | 41053 | 3 | 65623 | 32872 | 4 | 73011 | 39562 | 4 |
| **Fla1_A** | - | - | - | - | - | - | - | - | - | - | - | - |
| **Fla1_B** | - | - | - | - | - | - | 37551 | 15762 | 1 | - | - | - |
| **Fla2_A** | - | - | - | - | - | - | - | - | - | - | - | - |
| **Fla2_B** | - | - | - | - | - | - | - | - | - | - | - | - |
| **Igd1_A** | - | - | - | - | - | - | - | - | - | - | - | - |
| **Igd1_B** | - | - | - | - | - | - | - | - | - | - | - | - |
| **Igd2_A** | - | - | - | - | - | - | - | - | - | - | - | - |
| **Igd2_B** | - | - | - | - | - | - | - | - | - | - | - | - |
| **Iti1_A** | - | - | - | - | - | - | - | - | - | - | - | - |
| **Iti1_B** | - | - | - | - | - | - | 53334 | 11290 | 1 | 13705 | 3924 | 1 |
| **Iti2_A** | - | - | - | - | - | - | - | - | - | - | - | - |
| **Iti2_B** | - | - | - | - | - | - | - | - | - | - | - | - |
| **Itv1_A** | - | - | - | - | - | - | - | - | - | - | - | - |
| **Itv1_B** | - | - | - | - | - | - | - | - | - | - | - | - |
| **Nip1_A** | - | - | - | - | - | - | 1936 | 361 | 2 | - | - | - |
| **Nip1_B** | - | - | - | - | - | - | - | - | - | - | - | - |
| **Nip2_A** | - | - | - | - | - | - | 14170 | 3198 | 3 | 12962 | 3265 | 2 |
| **Nip2_B** | 12754 | - | - | - | - | - | 16235 | 3813 | 1 | - | - | - |
| **Nip3_A** | - | - | - | - | - | - | - | - | - | 14951 | 3355 | 3 |
| **Nip3_B** | - | - | - | - | - | - | 3088 | 1185 | 2 | - | - | - |
| **Nuu1_A** | - | - | - | - | - | - | - | - | - | - | - | - |
| **Nuu1_B** | - | - | - | - | - | - | - | - | - | - | - | - |
| **Nuu2_A** | - | - | - | - | - | - | - | - | - | - | - | - |
| **Nuu2_B** | - | - | - | - | - | - | - | - | - | - | - | - |
| **Qaj1_A** | - | - | - | - | - | - | - | - | - | 111 | - | - |
| **Qaj1_B** | - | - | - | - | - | - | - | - | - | - | - | - |
| **Qaj2_A** | - | - | - | - | - | - | - | - | - | - | - | - |
| **Qaj2_B** | - | - | - | - | - | - | - | - | - | - | - | - |
| **Qeq1_A** | - | - | - | - | - | - | - | - | - | - | - | - |
| **Qeq1_B** | - | - | - | - | - | - | - | - | - | - | - | - |
| **Qeq2_A** | - | - | - | - | - | - | - | - | - | 4844 | - | - |
| **Qeq2_B** | - | - | - | - | - | - | - | - | - | - | - | - |
| **V511_A** | - | - | - | - | - | - | - | - | - | - | - | - |
| **V511_B** | - | - | - | - | - | - | - | - | - | - | - | - |
| **V512_A** | 10433 | - | - | 83039 | 37 | 1 | - | - | - | - | - | - |
| **V512_B** | - | - | - | 10913 | - | - | 38823 | 14637 | 1 | - | - | - |
| **V541_A** | - | - | - | 2191 | 746 | 1 | 20436 | 2611 | 3 | 35897 | 2820 | 3 |
| **V541_B** | - | - | - | - | - | - | - | - | - | - | - | - |
| **V542_A** | - | - | - | - | - | - | - | - | - | - | - | - |
| **V542_B** | - | - | - | 164765 | 58663 | 1 | - | - | - | - | - | - |

### Table 6. Sequencing information, controls.

| Sample | 12Sv5 | | | Mam16S | | | Fish16S | | | 12SAH | | |
|---|---|---|---|---|---|---|---|---|---|---|---|---|
| | Raw count | Filt. count | Uniq. count | Raw count | Filt. count | Uniq. count | Raw count | Filt. count | Uniq. count | Raw count | Filt. count | Uniq. count |
| **EB1_I** | 9538 | 6422 | 1 | - | - | - | - | - | - | 3037 | - | - |
| **EB1_II** | 6788 | 5953 | 1 | - | - | - | - | - | - | - | - | - |
| **EB2_I** | - | - | - | - | - | - | - | - | - | - | - | - |
| **EB2_II** | 282 | 214 | 1 | 266 | 30 | 1 | - | - | - | - | - | - |
| **EB3_I** | 16 | - | - | 7 | - | - | - | - | - | - | - | - |
| **EB3_II** | - | - | - | - | - | - | 8 | - | - | - | - | - |
| **EB4** | 77014 | 60574 | 1 | 154 | - | - | - | - | - | 6 | - | - |
| **EB5** | 1 | - | - | 818 | 25 | 1 | - | - | - | 7 | - | - |
| **EB6** | 1 | - | - | 58220 | - | - | - | - | - | 53 | - | - |
| **EB7** | 3 | - | - | - | - | - | - | - | - | - | - | - |
| **GB1_I** | 10724 | 9378 | 2 | - | - | - | - | - | - | - | - | - |
| **GB1_II** | 11799 | 8077 | 1 | - | - | - | - | - | - | - | - | - |
| **GB2_I** | - | - | - | - | - | - | - | - | - | - | - | - |
| **GB2_II** | - | - | - | - | - | - | - | - | - | - | - | - |
| **GB3_A** | 2 | - | - | 542 | - | - | - | - | - | - | - | - |
| **GB3_B** | 7 | - | - | - | - | - | - | - | - | - | - | - |
| **GB4_A** | 6 | - | - | - | - | - | - | - | - | - | - | - |
| **GB4_B** | 67 | - | - | 347556 | 293445 | 1 | - | - | - | - | - | - |
| **PB1** | 49 | - | - | - | - | - | - | - | - | - | - | - |
| **PB2** | - | - | - | - | - | - | - | - | - | - | - | - |
| **PB3** | - | - | - | 6360 | 11 | 1 | - | - | - | - | - | - |
| **PB4** | 43460 | 30601 | 1 | 1737 | - | - | - | - | - | - | - | - |
| **PB5** | 15 | - | - | - | - | - | 5 | - | - | - | - | - |
| **PB6** | 2169 | 1492 | 3 | - | - | - | 16150 | - | - | 260 | - | - |
| **PB7** | 70951 | 54517 | 1 | - | - | - | - | - | - | - | - | - |
| **PB8** | 605 | - | - | - | - | - | - | - | - | - | - | - |

**Table 7. Species identified.** *Identified with shotgun metagenomics exclusively. $Identified with the targeted whale assay exclusively.

| Taxon | Common name | Found before in studied sites? | Sites (n=11) | Samples (50 bones) (n=46 samples) |
|---|---|---|---|---|
| Gadus morhua/macrocephalus/ogac | Cod | yes | 2 | 8 |
| *Mallotus villosus* | Capelin | yes | 3 | 4 |
| *Anarhichas lupus* | Atlantic wolffish | no | 2 | 7 |
| *Myoxocephalus scorpius* | Shorthorn sculpin | yes | 2 | 6 |
| *Cyclopterus lumpus* | Lumpfish | no | 1 | 1 |
| *Sebastes* | Redfish | no | 1 | 1 |
| *Hippoglossoides platessoides* | American plaice | no | 2 | 2 |
| *Hippoglossus hippoglossus* | Atlantic halibut | yes | 1 | 1 |
| *Salvelinus alpinus* | Arctic char | yes | 1 | 1 |
| *Alca torda* | Razorbill | yes | 1 | 1 |
| *Alle alle* | Little auk | yes | 1 | 1 |
| *Pinguinus impennis** | Great auk | yes | 2 | 4 |
| *Uria lomvia* | Brünnich's guillemot | yes | 4 | 8 |
| *Larus* | Gull | yes | 7 | 20 |
| Corvus | Raven | yes | 2 | 2 |
| *Phalacrocorax carbo* | Great cormorant | yes | 2 | 2 |
| *Mergus/Somateria* | Eider/merganser | yes | 6 | 12 |
| *Anas* | Dabbling ducks | yes | 1 | 1 |
| *Mergus* | Merganser | yes | 1 | 2 |
| *Somateria* | Eider | yes | 3 | 7 |
| Anserinae | Geese and swans | yes | 4 | 8 |
| *Anser* | Grey Geese | yes | 1 | 1 |
| *Branta* | Black Geese | yes | 1 | 2 |
| *Lagopus muta* | Ptarmigan | yes | 5 | 14 |
| *Lepus arcticus* | Arctic hare | yes | 3 | 6 |
| *Dicrostonyx groenlandicus* | Northern collared lemming | no | 1 | 4 |
| *Bos taurus* | Cattle | yes | 2 | 8 |
| *Capra hircus* | Goat | yes | 2 | 8 |
| *Ovibos moschatus* | Muskox | yes | 2 | 2 |
| *Ovis* | Sheep | yes | 2 | 8 |
| *Rangifer tarandus* | Caribou | yes | 7 | 29 |
| *Balaena mysticetus*$ | Bowhead whale | yes | 6 | 7 |
| *Balaenoptera physalus* | Fin whale | no | 1 | 1 |
| *Monodon monoceros* | Narwhal | yes | 2 | 7 |
| *Phocoena phocoena* | Harbour porpoise | yes | 1 | 2 |
| *Physeter catodon* | Sperm whale | yes | 1 | 1 |
| *Canis lupus* | Dog/wolf | yes | 9 | 19 |
| *Vulpes lagopus* | Arctic fox | yes | 9 | 16 |
| *Odobenus rosmarus* | Walrus | yes | 2 | 5 |
| *Erignathus barbatus* | Bearded seal | yes | 3 | 6 |
| *Pagophilus groenlandicus* | Harp seal | yes | 10 | 27 |
| *Phoca vitulina* | Harbor seal | yes | 5 | 14 |
| *Pusa hispida* | Ringed seal | yes | 7 | 15 |
| *Ursus maritimus* | Polar bear | yes | 1 | 2 |

**Table 8. Species identified per site using BBM.** *New species detection at the site as compared to morphological identifications. References for morphological identifications are reported in Table 2. $Potential contamination.

| | Fla | Nuu | Iff | Qoo | Igd | Iti | Nip | Qaj | Qeq | V51 | V54 |
|---|---|---|---|---|---|---|---|---|---|---|---|
| Gadus morhua/macrocephalus/ogac | - | - | 4 | 4 | - | - | - | - | - | - | - |
| *Mallotus villosus* | - | - | 2 | 1 | - | - | - | - | - | 1* | - |
| *Anarhichas lupus* | - | - | 2 | 5 | - | - | - | - | - | - | - |
| *Myoxocephalus scorpius* | - | - | 3 | 3 | - | - | - | - | - | - | - |
| *Cyclopterus lumpus* | - | - | - | 1 | - | - | - | - | - | - | - |
| *Sebastes* | - | - | 1 | - | - | - | - | - | - | - | - |
| *Hippoglossoides platessoides* | - | - | 1 | 1 | - | - | - | - | - | - | - |
| *Hippoglossus hippoglossus* | - | - | - | - | - | - | 1* | - | - | - | - |
| *Salvelinus alpinus* | - | - | - | - | - | - | - | - | - | - | 1 |
| *Alca torda* | - | - | - | - | - | - | 1 | - | - | - | - |
| *Alle alle* | - | - | - | - | - | - | - | - | 1 | - | - |
| *Pinguinus impennis* | - | - | 1 | 3 | - | - | - | - | - | - | - |
| *Uria lomvia* | - | - | 3 | 3 | 1 | - | - | 1 | - | - | - |
| *Larus* | - | 1 | 4 | 5 | - | - | 5 | 2 | 2 | - | 1* |
| Corvus | - | - | - | - | - | - | - | - | 1 | 1* | - |
| *Phalacrocorax carbo* | - | - | - | - | - | - | 1 | - | - | - | 1* |
| *Mergus/Somateria* | - | - | - | 3 | 1 | 2 | 4 | 1 | - | - | 1* |
| *Anas* | - | - | - | 1 | - | - | - | - | - | - | - |
| *Mergus* | - | - | - | 2 | - | - | - | - | - | - | - |
| *Somateria* | - | - | - | 3 | - | 1 | 3 | - | - | - | - |
| Anserinae | 3 | - | - | - | - | 1* | 3 | - | - | - | 1 |
| *Anser* | 1 | - | - | - | - | - | - | - | - | - | - |
| *Branta* | - | - | - | - | - | - | 2 | - | - | - | - |
| *Lagopus muta* | - | - | 4 | 4 | - | - | 2 | - | - | 2* | 2 |
| *Lepus arcticus* | 4 | - | - | - | - | - | 1 | - | - | - | 1 |
| *Dicrostonyx groenlandicus* | 4 | - | - | - | - | - | - | - | - | - | - |
| *Bos taurus* | - | - | - | - | - | 2$ | 1$ | - | - | 4 | 4 |
| *Capra hircus* | - | - | - | - | - | - | - | - | - | 4 | 4 |
| *Ovis* | - | - | 1$ | 1$ | - | - | - | - | - | 3 | 4 |
| *Rangifer tarandus* | 4 | - | 4 | 4 | - | 3 | 6 | - | - | 4 | 4 |
| *Balaena mysticetus* | 1 | 2 | 1 | - | - | - | - | - | - | 1* | 1* |
| *Balaenoptera physalus* | - | - | 1 | - | - | - | - | - | - | - | - |
| *Monodon monoceros* | 4 | 3 | - | - | - | - | - | - | - | - | - |
| *Phocoena phocoena* | - | - | - | - | - | - | 2 | - | - | - | - |
| *Physeter catodon* | - | - | - | - | - | - | - | - | - | 1 | - |
| *Canis lupus* | 3 | 1 | 2 | 3 | 1 | - | - | 2 | 1 | 2 | 4 |
| *Vulpes lagopus* | 1 | 1 | 3 | 3 | 1 | - | 1 | 1 | 4 | - | 1* |
| *Odobenus rosmarus* | - | 1 | - | - | - | - | - | - | - | 4 | - |
| *Erignathus barbatus* | 4 | 1 | - | - | - | - | 1 | - | - | - | - |
| *Pagophilus groenlandicus* | 2 | - | 4 | 6 | 2 | 1 | 1 | 2 | 4 | 2 | 3 |
| *Phoca vitulina* | - | - | - | - | - | 1* | 6 | - | 1 | 3 | 3 |
| *Pusa hispida* | 4 | 3 | - | 1 | - | - | 1 | 2 | 3 | - | 1* |
| *Ursus maritimus* | - | 2 | - | - | - | - | - | - | - | - | - |

**Table 9. Species identified with shotgun metagenomics at Iffiartarfik and Qoornoq.** Read counts of all taxa identified by the lowest common ancestor approach. Note that one species can be identified at several taxonomic levels. Dendrobatidae represents a false positive identified due to a short repetitive sequence in in the control region of two mitochondrial references genomes from the family Dendrobatidae.

| Raw assigned node | Metabarcoding equivalent | If1_A | If2_A | Qo1_A | Qo2_A | Qo3_A |
|---|---|---|---|---|---|---|
| Gadidae | *Gadus morhua/macrocephalus/ogac* | 36 | 9 | 42 | - | 75 |
| *Gadus macrocephalus* | *Gadus morhua/macrocephalus/ogac* | 5 | 1 | 5 | - | 14 |
| *Gadus ogac* | *Gadus morhua/macrocephalus/ogac* | 12 | 1 | 8 | - | 30 |
| *Gadus* | *Gadus morhua/macrocephalus/ogac* | 52 | 26 | 84 | - | 111 |
| *Anarhichadidae* | *Anarhichas lupus* | 2 | - | 2 | - | 8 |
| *Anarhichas lupus* | *Anarhichas lupus* | 13 | - | 38 | - | 90 |
| *Anarhichas* | *Anarhichas lupus* | 15 | - | 20 | - | 69 |
| *Cottidae* | *Myoxocephalus scorpius* | 13 | 5 | - | - | - |
| *Myoxocephalus scorpius* | *Myoxocephalus scorpius* | 80 | 33 | - | 6 | - |
| *Myoxocephalus* | *Myoxocephalus scorpius* | 12 | 5 | - | 1 | - |
| *Pinguinus impennis* | - | 1 | 2 | 2 | 4 | 9 |
| *Laridae* | *Larus* | 7 | 91 | 1 | 13 | 127 |
| *Larus dominicanus* | *Larus* | 14 | 191 | 1 | 38 | 284 |
| *Larus* | *Larus* | 6 | 78 | - | 10 | 94 |
| *Larus crassirostris* | *Larus* | - | 16 | 1 | 6 | 23 |
| *Lagopus muta* | *Lagopus muta* | 548 | 479 | 11 | - | 352 |
| *Lagopus* | *Lagopus muta* | 69 | 79 | 1 | - | 48 |
| *Phasianidae* | *Lagopus muta* | 47 | 50 | 2 | 1 | 35 |
| *Tetraoninae* | *Lagopus muta* | 44 | 29 | 2 | - | 22 |
| *Rangifer tarandus* | *Rangifer tarandus* | 10 | 8 | - | - | 1 |
| *Canis lupus* | *Canis lupus* | - | - | - | 22 | - |
| *Vulpes lagopus* | *Vulpes lagopus* | - | 3 | 18 | - | - |
| *Phoca* | *Pagophilus groenlandicus* | 19 | 24 | 7 | 13 | 14 |
| *Phocidae* | *Pagophilusgroenlandicus/Pusa hispida* | 67 | 88 | 60 | 60 | 54 |
| *Pagophilus groenlandicus* | *Pagophilus groenlandicus* | 401 | 386 | 76 | 176 | 294 |
| *Pusa hispida* | *Pusa hispida* | 2 | 5 | 14 | 22 | - |
| *Pusa* | *Pusa hispida* | - | 1 | 10 | 6 | 2 |
| *Dendrobatidae* | FALSE POSITIVE | 8 | 4 | 3 | - | 2 |

### Supplementary Note 1 - A comparison between metabarcoding and shotgun metagenomics

Our comparison of metabarcoding and shotgun metagenomics on DNA extracted from bulk bone highlights some of the commonly discussed pros and cons for each method[1,18]. We compared the performance of both methods by sequencing shotgun libraries from 5 samples at two sites (Iffiartarfik; 2 samples and Qoornoq; 3 samples) and comparing these with metabarcoding data from the same sites (Supplementary Figure 1). We found that, despite sequencing 1/6 the number of reads (metabarcoding: 11.5 million reads, shotgun metagenomics: 74.2 million reads), metabarcoding detected a far greater species diversity (21 species) than shotgun metagenomics (11 species; Supplementary Figure 1a). Furthermore, all but one of the species (the great auk; *Pinguinus impennis*) detected by shotgun metagenomics was also detected by metabarcoding. This finding demonstrates the high sensitivity of PCR in combination with specific primers utilised in metabarcoding. However, the sensitivity comes with a cost as the metabarcoding approach also detects contaminant sequences from sheep (*Ovis aries*). Furthermore, one species, the great auk (*Pinguinus impennis*), was detected by shotgun metagenomics, but not by metabarcoding. The apparent absence of great auk DNA in the metabarcoding samples most likely reflects that the bird assays used here (*12SAH* & *12SV5*), have a mismatch towards the 3' end of the reverse primer when compared to the great auk mitochondrial genome (NC_031347). This dissimilarity could prevent amplification of great auk DNA. Hence, these results highlight one of the advantages of shotgun metagenomics over metabarcoding: that the shotgun DNA library is built without amplification with specific primers, which bypasses the biases that this step might introduce.

The main advantage most often highlighted for shotgun metagenomics over metabarcoding is the accurate representation of abundances between species in a sample[19]. For metabarcoding experiments that utilises multiple assays, it is impossible to infer meaningful relations of the abundances between species detected by different assays. Furthermore, even taxa detected by the same assay can have heavily skewed abundances because of stochasticity between samples and primer binding bias[20]. As a consequence, it is common practice to convert metabarcoding data from read counts to presence/absence data.

We found that our metabarcoding data reflects the general abundance pattern of the shotgun metagenomics data. This is illustrated by estimates of relative abundance from metabarcoding and shotgun sequencing which form significant linear relationships (Supplementary Figure 2 and 3). Hence, although the metabarcoding approach is more biased than shotgun metagenomics, read counts in metabarcoding data can be used as a proxy for DNA abundance.

Lastly, we assessed how bulk bone performed as a substrate for shotgun metagenomic analyses by comparing our shotgun metagenomic data from bulk bone with similar shotgun data extracted from sediment in Seersholm et al. 2016[1]. We found that the bulk bone samples contain on average 23 times as much vertebrate DNA per megaread sequenced as the sediment samples do (Supplementary Figure 1b). Surprisingly, the amount of plant DNA is not higher in the sediment samples than in the bulk bone samples. These observations indicate that: (1) By using bulk bone as a substrate instead of sediment, it's possible to reduce the amount of sequencing by ~20 times, and still obtain similar numbers of vertebrate reads. (2) Bulk bone can also provide information on the surrounding vegetation. (3) The low amounts of vertebrate DNA in a typical sediment sample is not explained by a high concentration of plant DNA. Instead, it is likely that the abundance of bacteria in sediment dilutes the vertebrate DNA considerably.

From a technical point of view, the findings in this paper adds tangible data to a long discussion on the pros and cons for both shotgun metagenomics and metabarcoding on environmental DNA samples. For the purposes of this study, metabarcoding outperforms shotgun metagenomics because of the increased species diversity detected with metabarcoding. Despite the lower species diversity detected with shotgun metagenomics, there is a long list of advantages with this approach: (1) Better abundance estimates, which is particularly relevant in studies of plant communities in sediment samples. (2) The ability to estimate ancient DNA damage patterns. (3) The opportunity to extract full mitochondrial or plastid genomes of the most abundant taxa. (4) The possibility to assess diversity across the tree of life in a sample, without the need to conduct further laboratory work. Still, as we analyse semi-synthetic bulk bone samples in this study abundance estimates are not relevant, and the increased species diversity afforded by metabarcoding combined with its lower cost makes this approach the most suitable.

### Supplementary Note 2 – Contamination issues

We identify three types of contamination in this study, all three are at low levels compared to the endogenous DNA in the test samples and can easily be identified and excluded from downstream analyses.

**Cross contamination**

First and most importantly, we identify low level cross contamination in samples with low amounts of endogenous DNA. This is exemplified by the rare detection of some of the most abundant species from the bulk bone samples in extraction and non-template PCR controls (*Pagophilus groenlandicus* and *Larus* sp. in PCR blank 6, “PB6” and Rangifer tarandus in extraction blank 5, “EB5”). This issue is a testament to the absence of amplifiable DNA in blank controls in combination with the very high number of control samples included in this sample (2 out of 26 blank controls had evidence of cross contamination). Furthermore, in one sample from Nipisat (Nip2_B2) we detect cross contamination from the Fladstrand samples. This cross contamination most likely arises from sample “Fla1_B1” which was located directly adjacent to “Nip2_B2” on the 96 well plate under PCR amplification. We identify narwhal (*Monodon monoceros*) and muskox (*Ovibos moschatus*) in the sample from Nipisat, both of which are detected in the Fladstrand sample genetically and have been identified previously by morphology at the site. None of these species have been identified before at Nipisat, and no ancient muskox remains have been identified in all of West and South Greenland. The species was introduced from Northeast Greenland to West Greenland in the years 1962 and 1965[21]. Furthermore, these species where not detected in sample Nip2_B1, which is a separate extraction of the same bulk bone pool as Nip2_B2. Hence, we conclude that this is a single instance of cross contamination, and we have excluded sample Nip2_B2 from the dataset.

**Background laboratory contamination**

Another type of contamination that we identify originates from background laboratory contamination. In our samples this is illustrated by the detection of pig (*Sus scrofa*) and human (*Homo sapiens*) in both test and blank controls. As no Greenlandic species have been identified previously as common laboratory contaminants this instance of contamination does not pose a problem to this study, and accordingly DNA from human and pig was excluded from downstream analyses.

**Contamination of bone material from the environment or from storage**

Lastly, we detect low amounts of DNA from two domestic species (*Bos* and *Ovis*) at one Paleo-Inuit and three Inuit sites (Nipisat, Iffiartarfik, Itinnera, and Qoornoq). These species are common in the area surrounding the Thule sites today, and it is likely that DNA from these modern species have contaminated the ancient remains by leeching of urine, feces and other organic matter from the surface and into the ground. Nevertheless, for two of these sites, which were used until historic times (Qoornoq and Iffiartarfik) it cannot be ruled that the detection of cattle and sheep DNA in fact stems from ancient remains. Lastly, cattle and sheep have also been identified as background laboratory contaminants[22,23], and it is equally possible that the detection of these species can be attributed to background contamination, and not contamination from the environment. Hence, to be conservative, we have excluded DNA from cow and sheep from all Inuit sites from downstream analyses.

In addition, we identify one instance of presumed contamination that is not easily accounted for. We identify Asian elephant (*Elephas maximus*) in two samples from Itinnera. The identification represents a single ASV detected at read counts of 6301 and 583 for Iti1_A2 and Iti2_A2, respectively. The relatively high read counts, and the detection of the same ASV across two different samples suggests that this identification could represent endogenous DNA. To investigate this further, we resequenced the two samples using the same *16S* primers. From this experiment, we detected the ASV matching Asian elephant in Iti2_A2 again. Hence, we conclude that the detection of Asian elephant DNA from two Ittinnera samples represents endogenous DNA in the sample and not spurious contamination.

The presence of remains from Asian elephant in Greenland during Saqqaq times is, however, highly unlikely. Remains from other proboscideans such as wholly mammoth (*Mammuthus primigenius*) or the American mastodon (*Mammut americanum*) could perhaps have drifted to Greenland by sea. However, both of these species are present in the reference database and provide lower similarity scores to the ASV (*Mammuthus primigenius*: 98.92% & *Mammut americanum*: 93.55%) than the match to the Asian elephant (*Elephas maximus*: 100%). On the other hand, there is only a single base pair difference between the match to mammoth and Asian elephant, and it is not impossible that unsampled local variation in the mitochondrial genome of mammoth accounts for this difference. To get a high resolution identification, we attempted to amplify control region elephant DNA from these samples using primers specific to both elephant and mammoth from Brandt et. al (2012)[24]. However, no amplification was observed from any of the samples.

In conclusion, the detection of probescidian DNA from two Itinnera samples represents endogenous DNA in the sample and not spurious contamination. It is likely that the detected DNA represents wholly mammoth remains that drifted to Greenland by sea. However, it is also possible that the samples from Itinnera have been cross contaminated with elephant DNA during the long storage of these samples at the Zoological Museum in Copenhagen.
